# Systematic assessment of multi-gene predictors of pan-cancer cell line sensitivity to drugs exploiting gene expression data

**DOI:** 10.1101/095224

**Authors:** Linh C. Nguyen, Cuong C. Dang, Pedro J. Ballester

## Abstract

Selected gene mutations are routinely used to guide the selection of cancer drugs for a given patient tumour. Large pharmacogenomic data sets were introduced to discover more of these single-gene markers of drug sensitivity. Very recently, machine learning regression has been used to investigate how well cancer cell line sensitivity to drugs is predicted depending on the type of molecular profile. The latter has revealed that gene expression data is the most predictive profile in the pan-cancer setting. However, no study to date has exploited GDSC data to systematically compare the performance of machine learning models based on multi-gene expression data against that of widely-used single-gene markers based on genomics data.

Here we present this systematic comparison using Random Forest (RF) classifiers exploiting the expression levels of 13,321 genes and an average of 501 tested cell lines per drug. To account for time-dependent batch effects in IC_50_ measurements, we employ independent test sets generated with more recent GDSC data than that used to train the predictors and show that this is a more realistic validation than K-fold cross-validation. Across 127 GDSC drugs, our results show that the single-gene markers unveiled by the MANOVA analysis tend to achieve higher precision than these RF-based multi-gene models, at the cost of generally having a poor recall (i.e. correctly detecting only a small part of the cell lines sensitive to the drug). Regarding overall classification performance, about two thirds of the drugs are better predicted by multi-gene RF classifiers. Among the drugs with the most predictive of these models, we found pyrimethamine, sunitinib and 17-AAG.

## Introduction

Personalised approaches to the diagnosis and treatment of cancer is a strong current trend, often based on the analysis of tumour DNA^1^. Somatic DNA mutations can affect the abundance and function of a range of gene products, including those involved in the response of the tumour to anticancer therapy^2^. Therefore, the genomic profile of a tumour is usually valuable for predicting its sensitivity to a certain drug^3,4^. Thus, a number of studies have profiled tumours using single-nucleotide variants or copy-number alterations to use them as input features to predict which tumours will be sensitive to a given drug. In addition, transcriptomic data has also been proven to be an informative molecular profile^5^, as the expression levels of genes have led the identification of cancer subtypes, prognosis prediction and drug sensitivity prediction^6^.

Human-derived cancer cell lines, especially immortalised cancer cell lines, play an important role in preclinical research for the discovery of genomic markers of drug sensitivity ^5,7–9^. This type of tumour models permits experiments to be implemented quickly and with a relatively low cost^10,11^, unlike more patient-relevant models like *ex vivo* tumour cultures^12,13^ or patient-derived xenografts^14,15^ (in contrast to these advantages, cell lines have also well-known limitations that have to be kept in mind^10^). The molecular profiles of such cell lines are often used as input features for drug sensitivity prediction ^5,8^ via the development of both single-gene markers and other models like pharmacogenomics^16–18^, pharmacotranscriptomics^19–21^,multi-task learning ^16,17,22–25^ and QSAR ^26,27^. Recently, several consortia have generated large pharmacogenomic datasets, which consist of both molecular and drug sensitivity profiles of several hundreds of cancer cell lines, e.g. Genomics of Drug Sensitivity in Cancer (GDSC)^8^, Cancer Cell Line Encyclopedia (CCLE)^9^ and that profiling the panel of 60 cancer cell lines from the National Cancer Institute (NCI-60)^7^. Among them, GDSC data is currently offering the highest number of cell lines tested per drug^8^.

To date, predictive models based on GDSC data have been mostly restricted to single-gene markers of drug sensitivity^8^ (i.e. statistically significant drug-gene associations). However, multi-gene elastic net models have also been used for a related purpose, namely estimating the importance of somatic mutations in drug sensitivity prediction^8^. Some of us have also investigated the performance of multi-gene machine learning models exploiting GDSC data^16^. Nevertheless, as other efforts ^9,7,18^, we did not study how well multi-gene markers compare to single-gene markers. Such analysis is essential to understand the benefits of modelling multiple gene alterations. Very recently, machine learning models have been used to compare the predictive value of various molecular profiles in drug sensitivity modelling^5^, although without comparing such models to single-gene markers. An important outcome of this study was revealing that gene expression data was the most predictive molecular profile in the pan-cancer setting. Beyond this research area, multi-variate machine learning models are also starting to be advocated for genomic-based prediction of other complex phenotypic traits^28^.

In practice, it is entirely possible that models based on one feature (single-gene markers) are more predictive than those based on more than one feature (multi-variate classifiers). In part, this is due to the high-dimensionality of training data (here, the number of gene expression values is much higher than that of cell lines treated with the considered drug), which poses a challenge to classifiers. Furthermore, as cell line sensitivity to a drug depends on its molecular features, the performance of models exploiting different molecular profiles will be drug-dependent. Therefore, the key question is for which drugs are multivariate markers more predictive of cell line sensitivity than univariate markers. Very recently, this question has been finally investigated using large-scale GDSC data^5^, although there are several limitations in this analysis. First, this study considered LOBICO logic models with up to four features because searching for more complex models was not feasible with LOBICO^5^, but a drug can have many more than four informative gene alterations. Second, machine learning models were only used to establish which molecular profiles were more informative on average across all drugs. Hence, the performances of these models were not compared against those of single-gene markers (this was only done with logic models). Third, both logic model selection and its classification performance assessment were carried using the same data folds in the adopted cross-validation procedure. Therefore, these cross-validated results represent an overoptimistic performance assessment of LOBICO models.

Here we study the performance of machine learning exploiting gene expression profiles. In addition, we compare the performance of these multi-gene machine learning models to that of single-gene markers. For each drug, this analysis is conducted by selecting its best single-gene marker and its multi-gene model on a training set representing the data available at model selection time. Thereafter, we test both models in an unbiased manner using a time-stamped independent test set, i.e. data that was generated after the training data and not used for model building or selection. The advantages of using a time-stamped data partition instead of K-fold cross-validation are that this mimics a blind test, the same data is not used for both model selection and performance assessment (thus avoiding performance overestimation) and real-world issues like time-dependent batch effects^29^ are taken into account. On the other hand, since transcriptomic data has been found to be the most predictive in the pan-cancer setting^5^, our study focuses on the exploitation of transcriptomic data. In particular, the predictive performance of pan-cancer markers of drug sensitivity on an independent test set is most relevant to help to stratify patients for basket trials^30^,where patients with any type of cancer are included if their tumours are predicted to be sensitive to the investigated treatment. Another reason to limiting the scope to transcriptomic-based machine learning models is that models integrating data from multiple molecular profiling technologies would be less amenable for clinical implementation due to much higher requirements in cost, time and resources per patient. Therefore, there is a need to understand for which drugs models combining gene expression values provide better cell line sensitivity prediction than standard single-gene markers.

## Results and discussion

### Comparing single-gene markers and transcriptomic-based models on the same test sets

A single-gene marker is a classifier considering the mutational status of a given gene as its only independent variable (i.e. whether this gene is wild-type or mutated). As the gene used as marker arises from the analysis of which drug-gene associations are statistically significant based on the training data, external validation of such markers is not carried out. In this sense, machine learning represents a different culture^31^, where the validity of the predictor is only demonstrated if its prediction is better than random on a test set independent of the employed training set. In this study, we use the same test set to compare the performance of both single-gene markers and multi-gene transcriptomic-based RF models.

For each drug, there are two data sets generated with non-overlapping sets of cancer cell lines. The first one is the training set, which contains cell lines that were tested prior to the release of release 1 of the GDSC data, each with its IC_50_ values for the drug and its gene expression profile. The second set is the test set, including the new cell lines from release 5 (i.e. new data not included in the first release). The median logIC_50_ in μM units of all cell lines in the training set defines the sensitivity threshold for both the training set and the test set. The next step is evaluating the performance of both methods in both data sets by calculating the Matthews Correlation Coefficient (MCC), Precision (PR), Recall (RC) and F1-score (F1). The Methods section provides further details on performance evaluation.

Random Forest (RF)^32^ is a machine learning technique that works well on high-dimensional data^33^, including GDSC data^16^. Therefore, without making any claim about its optimality, we constructed a RF classification model on the same training data set as the single-gene marker. This permits a direct comparison of the two models. Each RF model was built using 1000 trees, with the default value of the control parameters m_try_ (the square root of the number of considered features). The built RF model is subsequently tested on the corresponding test set.

Figure 1 displays the results for the drug pyrimethamine as an example. Pyrimethamine targets DHFR in the DNA replication pathway^34^ and its strongest association is to the BRAF gene (P=0.002) leading to a moderate level of prediction on this training set (Figure 1A). The prediction of this single-gene marker on the test set (Figure 1B) is worse than random (MCC=0.03), with its recall being particularly poor (RC=0.03) and average precision (P=0.50).Unsurprisingly, RF prediction on the training set is perfect due to intense overfitting^35^ arising from the high dimensionality of the problem (Figure 1C). Nevertheless, it is important to note that this overfitted model achieves a substantially better test set performance than that of the best single-gene marker (compare Figures 1D and 1B, respectively).

**Figure 1.**
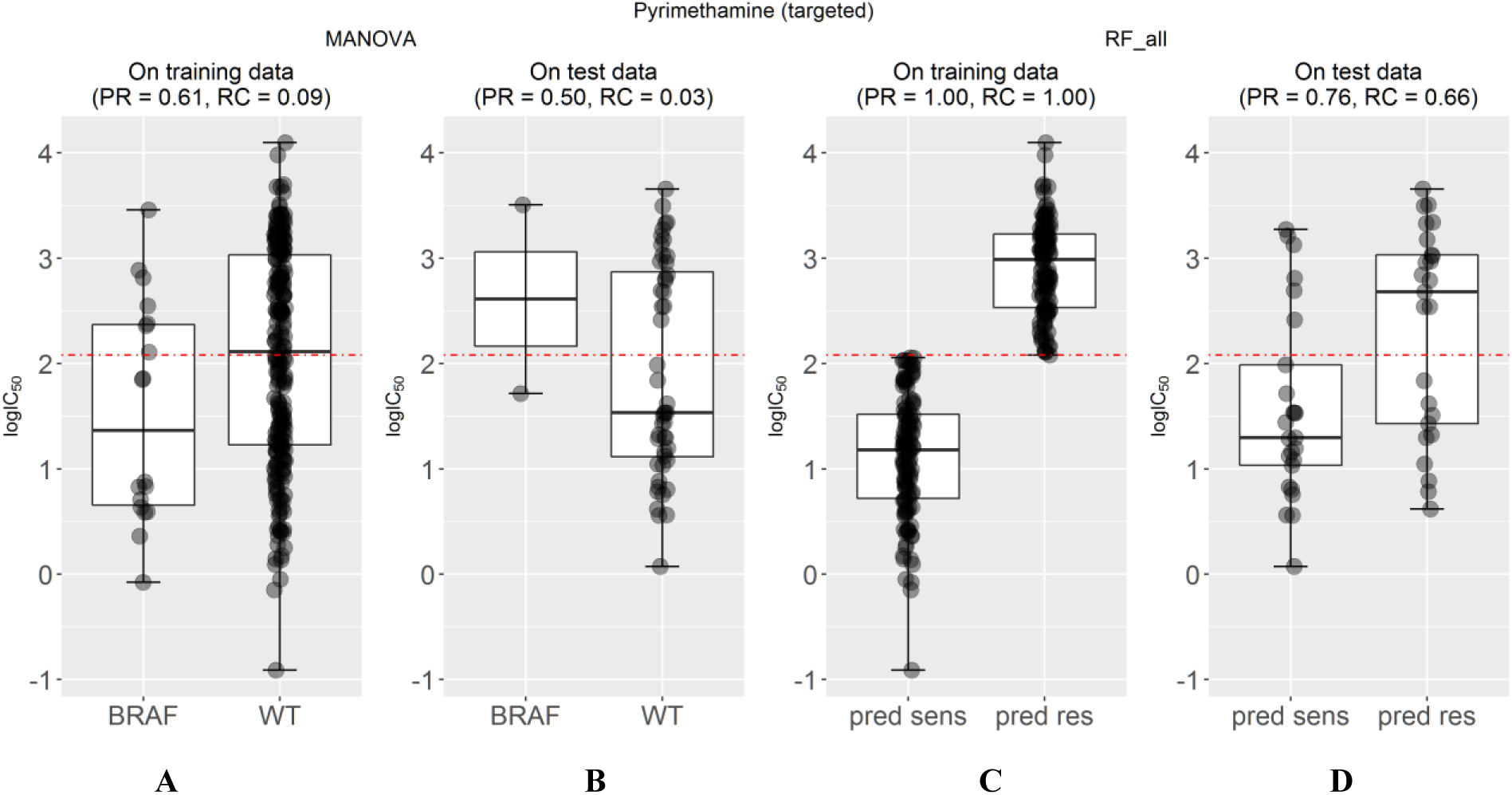
Predictive performance of markers of cell sensitivity to the approved drug pyrimethamine. **(A)** The single-gene marker with the lowest p-value on the training set was the pyrimethamine-BRAF sensitising association (P=0.002)^8^. **(B)** The boxplots show the sensitivity of cell lines on the independent test set for pyrimethamine depending on whether these harbour mutations in the BRAF gene or not (WT). Using this marker, BRAF-mutant cell lines are predicted to be sensitive to this drug (i.e. below the threshold in red established with training data), but the prediction is worse than random (MCC=-0.03) with its recall being particularly poor (RC=0.03) and average precision (P=0.50). **(C)** The multi-gene marker was built using RF and the gene expression profile on exactly the same drug-cell pairs as the single-gene markers. **(D)** On the test set, the RF classifier achieves a substantially better performance than single-gene markers (MCC=0.36 vs −0.03) with PR=0.76 and RC=0.66.

### Large inter-drug variability in the response rate of cell lines predicted to be sensitive

To assess the proportion of cell lines predicted to be sensitive that are actually sensitive to a drug by each model, we calculated their precision (PR) on the test set. Figure 2 shows the comparison between test set precision of single-gene markers and that of multi-gene models across 127 drugs. The precision of each method is highly drug-dependent and 61 drugs had its best single-gene marker leading to higher precision than the corresponding multi-gene model, whereas the other 66 drugs had the multi-gene model with better precision (see results.MNV.RF-CV.RF-TEST.xlsx in the Supplementary Information). In other words, the sensitivity of cancer cell lines against 66 drugs can be predicted with higher precision exploiting multi-variate gene expression data rather than a single gene mutation. In particular, the multi-gene model provides better precision for all the drugs for which the best single-gene marker involves a relatively rare mutation (i.e. those for which no test set cell line is mutated with respect to the marker gene and thus are unable to provide any level of precision).

**Figure 2:**
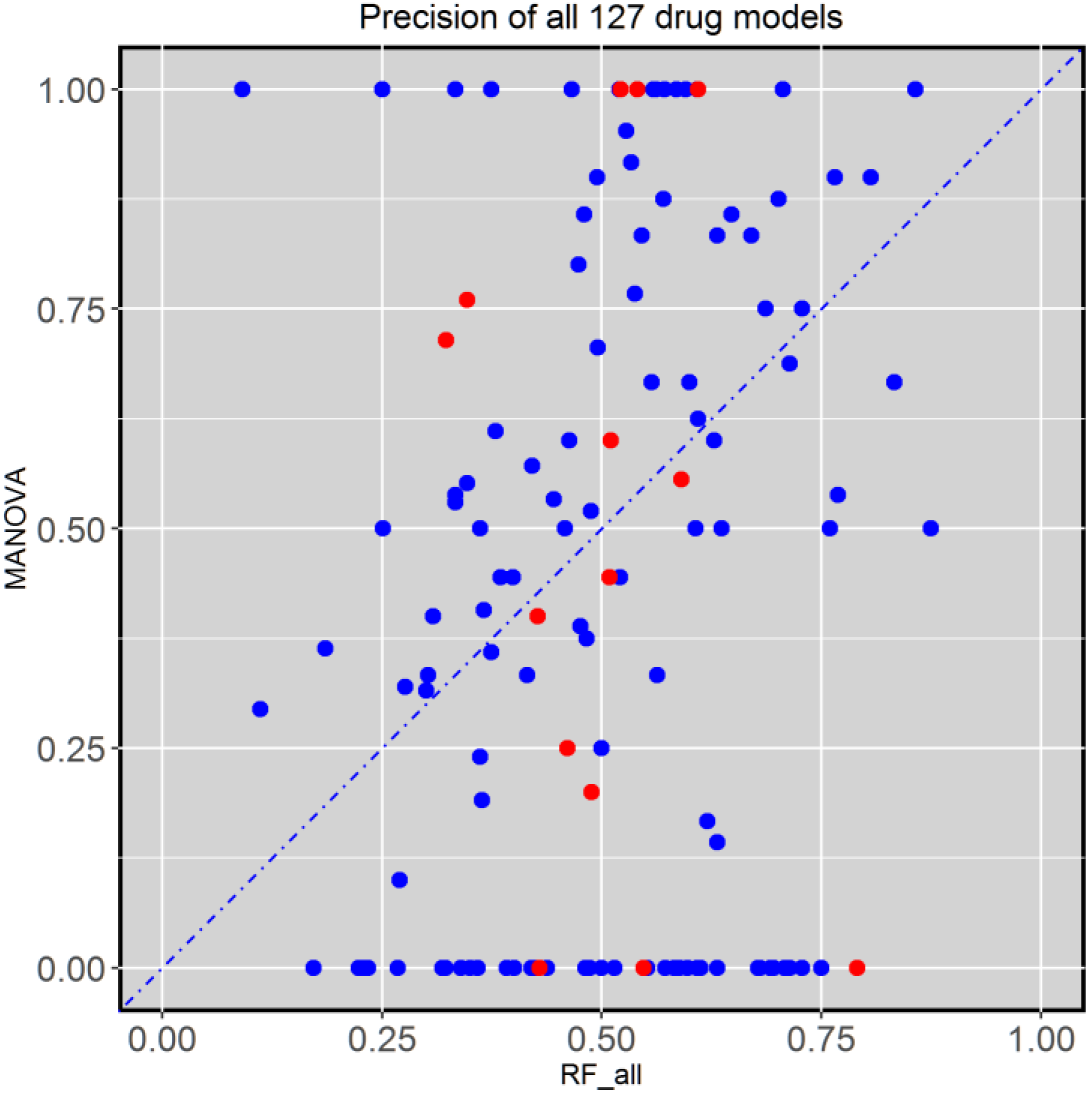
Test set precision of MANOVA single-gene markers versus RF transcriptomic models across the 127 drugs. A large variability is observed, with 66 drugs obtaining better precision with RF classifiers using all transcriptomic features. Cytotoxic drugs are red-coloured and targeted drugs are blue-coloured.

Next we present two examples of drugs for which the test set precision generated by the multi-gene model is higher than that of the single-gene model (Figure 3). AZD628 is a b-raf inhibitor, which plays a regulatory role in the MAPK/ERK pathway^36^. This drug is associated with the mutations in the BRAF gene (P=3·10^-15^), which codes for the b-raf kinase. 50% of BRAF-mutant cell lines are sensitive to this drug, while by using the RF model combining all 13,321 transcriptomic features, 88% of cell lines predicted to be sensitive are actually sensitive to this drug. The second example is the prediction of sensitivity to sunitinib, which targets multiple receptor tyrosine kinases regulating different aspects of cell signaling^37^. The most strongly associated gene to sunitinib is KDR (P=0.0002). As no KDR mutation was found in any test cell lines, the single-gene marker could not predict the sensitivity of any cell line to sunitinib (PR=0). In contrast, multi-gene model provides a much better precision for this drug (PR=0.66). The multi-gene models of both drugs generate a higher recall than their corresponding single-gene model, which is investigated in the next section.

**Figure 3:**
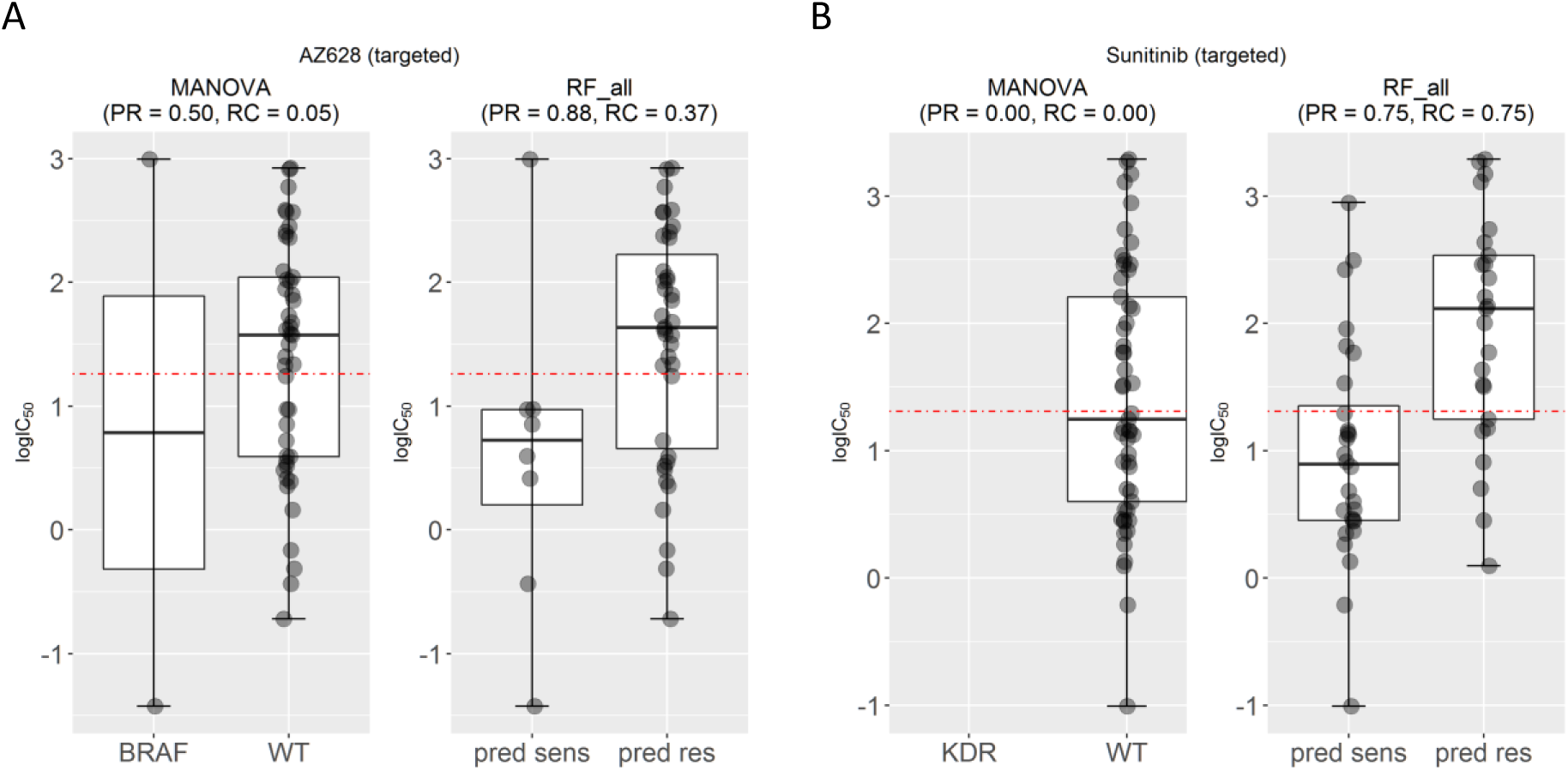
Examples of drugs for which transcriptomic markers predict their cell line sensitivity with better precision than that of single-gene markers on the test set. **(A)** Test set precision obtained by the AZD628-BRAF marker is moderate (PR=0.50) despite being a strong drug-gene association (P=3·10^-15^). By contrast, the multi-gene marker for AZD628 achieves a substantially higher precision (PR=0.88). **(B)** The Sunitinib-KDR association (P=0.0002) offers no precision in the test set because none of the test cell lines harbour mutations in the KDR gene. By contrast, the transcriptomic marker achieves a much higher precision (PR=0.75). Interestingly, both multi-gene markers achieve much better recall (RC=0.37 and RC=0.75) than their corresponding single-gene markers (RC=0.05 and RC=0.00), which means that a substantially higher proportion of sensitive cell lines are correctly predicted as sensitive.

### Multi-gene markers generally achieve much higher recall than single-gene markers

Figure 3 shows that the test set recall is much higher for multi-gene markers than for single-gene markers of AZD628 and sunitinib. To examine whether this is a general trend, Figure 4A plots test set recall across all the drugs. This is indeed a clear trend: 119 out of 127 drugs obtain a higher proportion of correctly predicted sensitive cell lines with the multi-gene markers.

Figure 4B shows the test set F-Score (F1) for the same drugs. High F1 values highlight markers achieving both high precision and high recall in the test set. Notably, the multi-gene classifiers lead to better recall and F1-scores in all the cytotoxic drugs. We have selected two drugs with high F1 by the multi-gene marker, which are BAY-61-3606 and 17-AAG in order to analyse them further (Figure 5).

**Figure 4:**
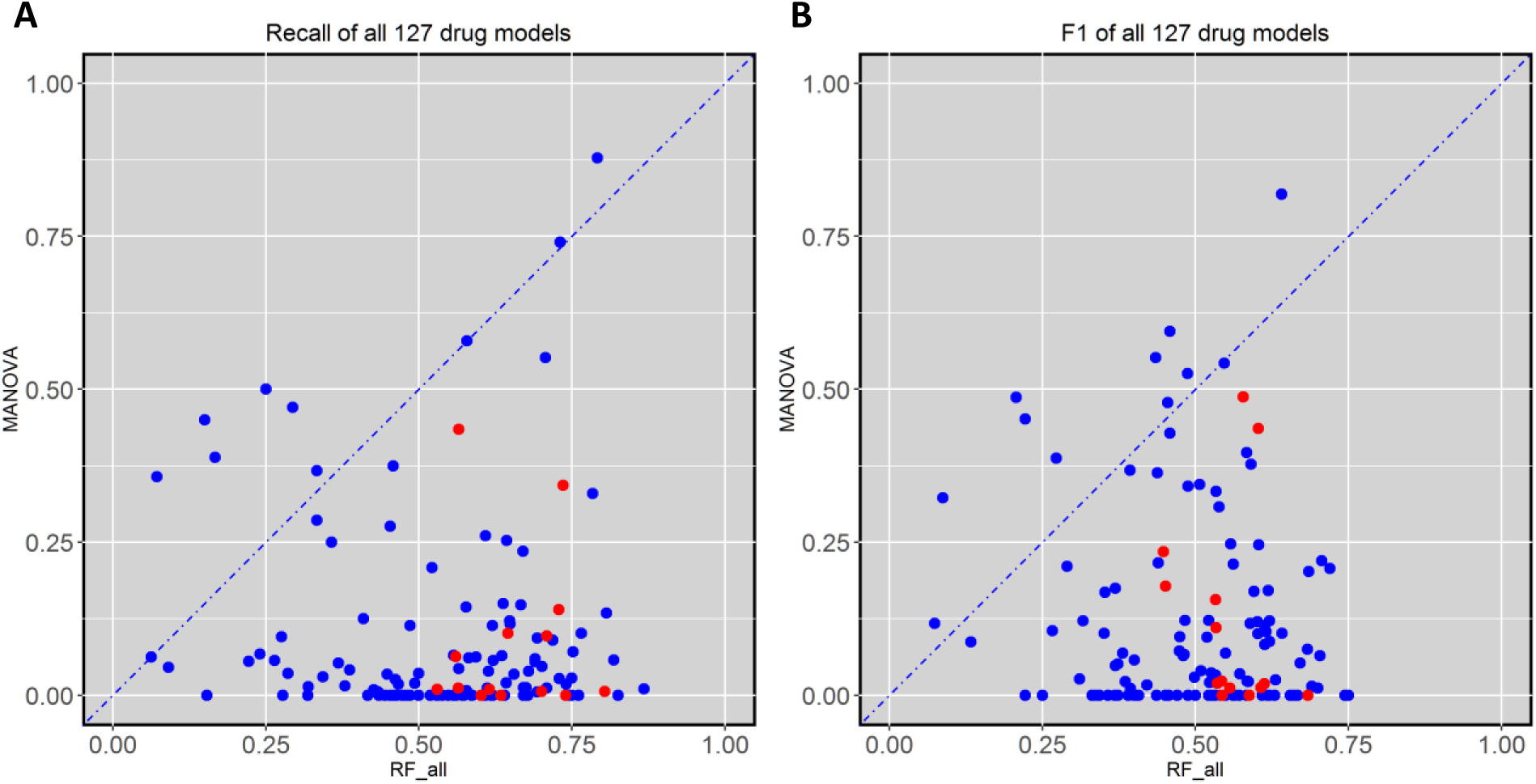
Test set recall and F-scores of single-gene and transcriptomic models across the 127 drugs. **(A)** transcriptomic markers achieve much higher recall than single-gene markers in 117 of the 127 drugs. **(B)** Similarly, multi-gene markers achieve higher F-scores in 117 of the 127 drugs. In each plot, cytotoxic drugs are red-coloured and targeted drugs are blue-coloured. All cytotoxic drugs have better recall and F-scores by the RF transcriptomic models.

**Figure 5:**
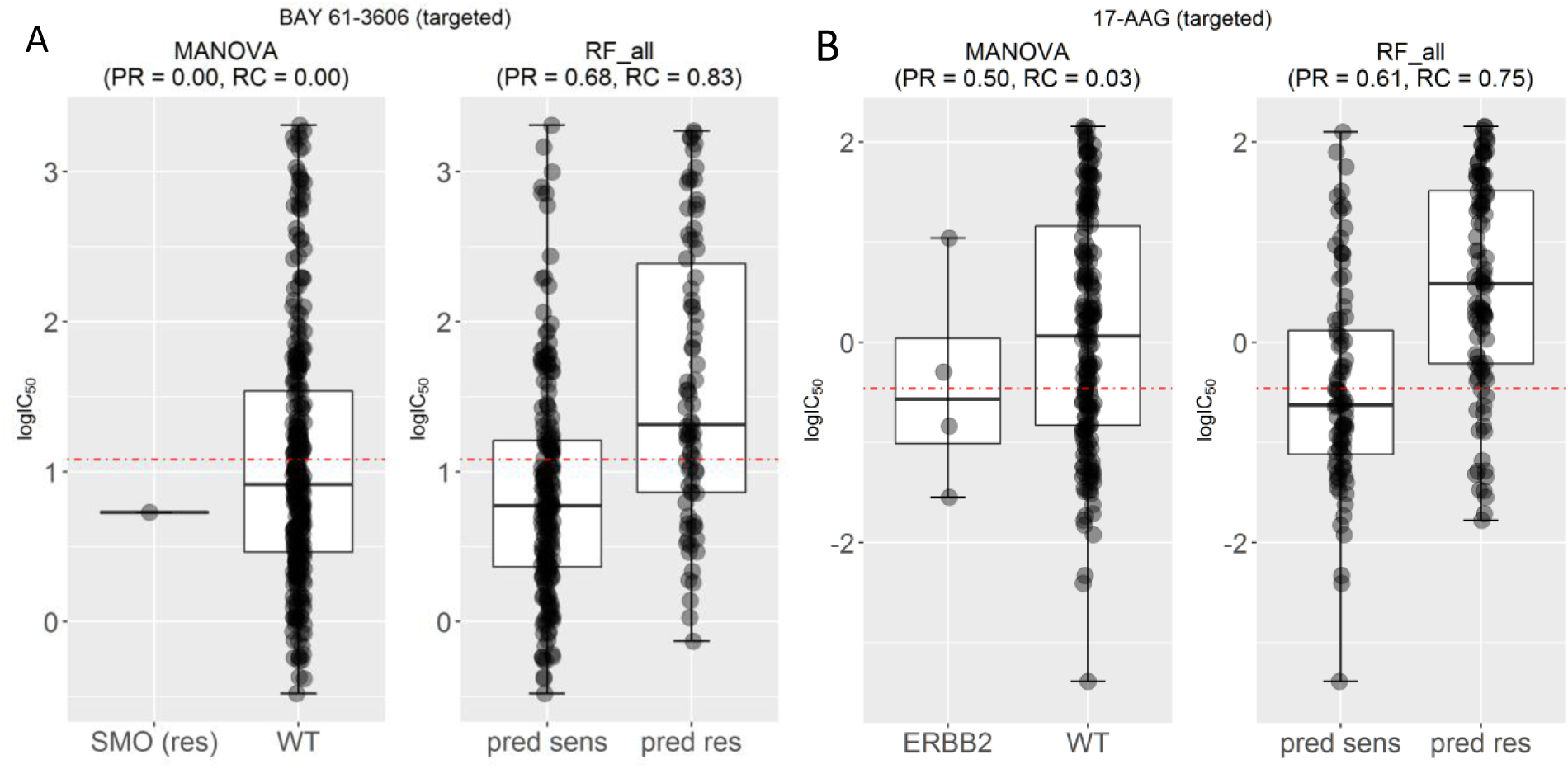
Examples of drugs that have high recall and F-scores. **(A)** Mutated SMO was the most significant single-gene marker for BAY-61-3606 resistance (P=0.03) using training data. On the test set, this marker obtained no precision and no recall because the only SMO-mutant test set cell line was misclassified. By contrast, the corresponding multi-gene marker, built with the same training data, obtained a high precision (PR=0.68) and better recall (RC=0.83) on the same test data. **(B)** Mutated ERBB2 is the most significant single-gene marker of 17-AAG sensitivity (P=0.008), but its test set recall is poor (RC=0.03). By contrast, the multi-gene marker achieves a much higher precision (PR=0.61) and recall (RC=0.75).

Figure 5A compare the test set performance between single-gene and multi-gene models for the drugs BAY-61-3606 and 17-AAG, respectively. BAY-61-3606 is an inhibitor for the spleen tyrosine kinase, with key roles in adaptive immune receptor signalling as well as the regulation cellular adhesion and vascular development^38^. The single-gene model generates poor precision and recall for this drug (PR = RC = 0), as the only cell line that harbour the actionable mutation was incorrectly predicted as resistant (TP = 0). By contrast, the multi-gene model achieves high performance in terms of both precision and recall (PR = 0.68 and RC = 0.83). On the other hand (Figure 5B), 17-AAG specifically inhibits HSP90, a protein that chaperones the folding of proteins required for tumour growth^39^. The multiple-gene model provides much higher PR (PR = 0.61) and RC (RC = 0.75) than with its best single-gene marker (PR = 0.50 and RC = 0.03). The latter exemplifies a common problem with single-gene markers: often only a small proportion of tumours harbour the actionable mutation^40^. This translates to very low recall, which in a clinical setting would mean that only a small part of the patients responsive to the drug would be treated with it because of being missed by its marker.

### The importance of using independent test sets to benchmark markers of drug sensitivity

After separately analysing the two sources of classification error via precision and recall, we analysed both types of error together in order to assess which predictors are better than random classification (i.e. MCC = 0)^41^.

The classification of both models can in principle be assessed in three ways across the considered drugs (Figure 6). Figure 6A evaluates the MCC of both predictors on the training data, which is common practice with single-gene markers. Figure 6C presents the evaluation of MCC on the non-overlapping test sets. Single-gene markers perform better on the training set than on the test set (on average, MCC_training_=0.11 vs MCC_test_=0.05; see Figures 6A and 6C), which is due at least in part to the identification of chance correlations in the training set. Unsurprisingly, multi-gene models perform much better on the training set due to intense overfitting (on average across drugs, MCC_training_=1 vs MCC_test_=0.12). However, despite overfitting, it is important to note that these models provide better average better test set performance than single-gene markers (MCC_test_=0.12 vs MCC_test_=0.05). This is a well-known characteristic of the RF technique, which is robust to overfitting in that it is able to provide competitive generalisation to other data sets despite overfitting (this behaviour has also been observed in analogous applications of RF^43^).

**Figure 6:**
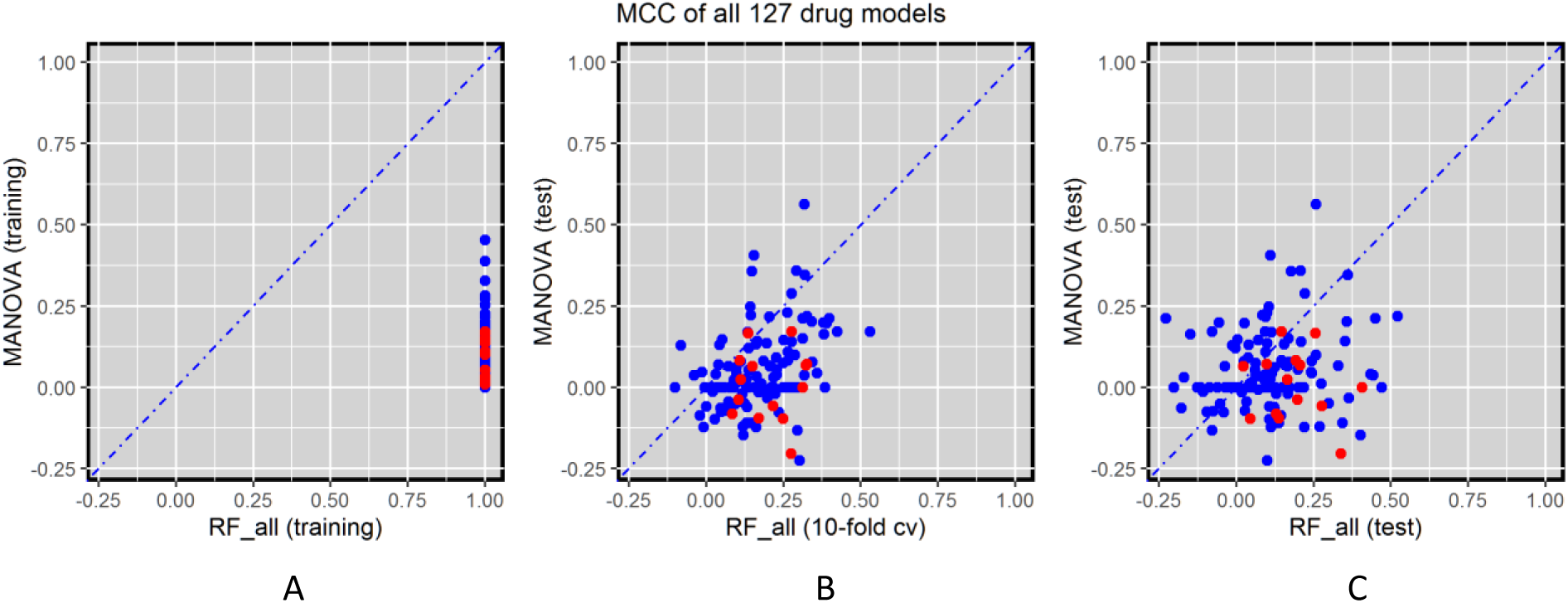
Global performance assessment of single-gene markers versus transcriptomic markers across the 127 drugs. **(A)** Performance assessment on the training data would be strongly biased towards multi-gene markers due to intense overfitting (given the high dimensionality of training data, multi-gene markers obtain maximum MCC for all drugs). **(B)** The performance of single-gene markers on the test set is compared to the 10-fold cross-validated performance of multi-gene markers using training data. The cross-validation is not used for model selection as there is only one RF model per drug (i.e. no RF control parameter is tuned because the recommended m_try_ is used due to the high dimensionality of each of the 127 classification problems). However, cross-validation results are substantially better than those from the test set with more recent GDSC data (MCC of 0.18 averaged over the drugs), which suggest time-dependent batch effects^10,42^. **(C)** Using all the comparable data released after the initial GDSC release as a time-stamped test set, 66.1% of drugs are better predicted by the transcriptomic features (this figure is 84.2% using cross-validation). This is the most realistic form of retrospective performance assessment, which leads to the worse results on this challenging problem (MCC of 0.12 averaged over the drugs).

Figure 6B shows the comparison between performance of the single-gene markers on the test set and the 10-fold cross-validated performance of the multi-gene markers on the training set. The latter provides a more optimistic performance assessment (average MCC=0.18 and 84.2% of drugs better predicted by the multi-gene models). This is likely due to effects between different batches of culture medium that are known to affect drug sensitivity measurements^10,42^. As expected, testing the models on the independent test sets generates worse results than on the training test or the cross-validation set.

## Conclusions

To the best of our knowledge, this is the first systematic comparison of single-gene markers versus transcriptomic-based machine learning models of cell line sensitivity to drugs. This is important as transcriptomic data has been shown to be the most predictive data type in the pan-cancer setting^5^. A closely related analysis was included in a very recent study^5^. However, this analysis is based on logic classifiers that can only exploit up to four features instead of fully-featured machine learning classifiers. Furthermore, the performance results in that study are based on cross-validations, thus leading to overoptimistic performance because of batch effects as we have seen here. The latter would be exacerbated if the same cross-validation is also used for model selection as it was the case in that study^5^. Despite these limitations, these new logic classifiers are very valuable in that they can potentially explain why a particular cell line is sensitive to the drug, something that machine learning classifiers are not suitable for.

Although single-gene markers were able to predict the sensitivity of cancer cell lines to anti-cancer drugs with generally high test set precision (Figure 2), very poor precision and very low recall was provided for other drugs, especially those that are best associated with relatively rare actionable mutation. On the other hand, multi-gene classifiers obtain much better recall, also known as sensitivity, for most of the drugs (Figure 4). This result is in line with criticism of single-gene markers, which lead to an extremely small proportion of patients that can benefit^40^. In this sense, one could argue that there is a need for not only precision oncology, but for precision and recall oncology, and that multi-variate classifiers have the potential to identify all the responsive patients, not only the subset of those with an actionable mutation.

While no strong single-gene markers of sensitivity were found for cytotoxic drugs^8^, the multi-gene machine learning models perform better than the single-gene markers in 12 of the 14 cytotoxic drugs (Figure 6C), with all cytotoxic drugs having better recall (Figure 4A). This suggests that the sensitivity to cytotoxic drugs has a stronger multi-factorial nature, which is thus better captured by multi-gene models. Although much less developed to date, personalised oncology approaches have already been suggested for cytotoxic drugs^44,45^.

The study of molecular markers for drug sensitivity is currently of great interest. This endeavour is not limited to improve personalised oncology, it is also important for drug development and clinical research^46,47^. As a part of cancer diagnosis and treatment research, a vast amount of tumour molecular profiling data is typically generated^48^ and thus there is an urgent need for their optimal exploitation^49^. Here we proposed a method to exploit transcriptomic data of cancer cell lines to classify them into sensitive and resistant groups. Our study has found that cancer cell sensitivity to two thirds of the studied drugs, including 12 of the 14 cytotoxic drugs, are better predicted with multi-variate transcriptomic-based RF classifiers. These models are particularly useful in those drugs where their best genomic markers are based on rare mutations. Another contribution of this study is in the proposal of a more realistic performance assessment of markers, which leads to less spectacular but more robust results. Beyond this proof-of-concept study across 127 drugs, there are several important avenues for future work, which are far too extensive to be incorporated here. For instance, there is a plethora of feature selection techniques that can be applied to reduce the dimensionality of the problem prior to training the classifier for a given drug. Furthermore, the predictive performance of these models can be evaluated on more data or integrated with other molecular profiles. Lastly, we have used a robust classifier technique, RF, but there are many other available and some of these must be more appropriate depending on the analysed drug.

## Methods

### GDSC pharmacogenomic data

From the Genomics of Drug Sensitivity in Cancer (GDSC) ftp server^50^, the following files from the first data release were downloaded: gdsc_manova_input_w1.csv and gdsc_manova_output_w1.csv. There are 130 unique drugs in gdsc_manova_input_w1.csv, as camptothecin was tested twice, drug IDs 195 and 1003, and thus we only kept the instance that was more widely tested (i.e. drug ID 1003 on 430 cell lines). Hence, the data represents a panel of 130 drugs tested against 638 cancer cell lines resulting in a total of 47748 IC_50_ values (57.6% of all possible drug-cell pairs). In addition, we downloaded new data from the latest release using the same experimental techniques to generate pharmacogenomic data and selected genes as in the first release (gdsc_manova_input_w5.csv). Release 5 contains 139 drugs tested on 708 cell lines comprising 79,401 IC_50_ values (80.7% of all possible drug-cell pairs). Hence, the majority of the new IC_50_ values come from previously untested drug-cell pairs formed by drugs and cell lines in common between both releases.

The downloaded “IC_50_” values are actually the natural logarithm of IC_50_ in μM units, so negative values come from drug responses more potent than 1μM. Each of these values were converted into their logarithm base 10 in μM units, denoted as logIC_50_ (e.g. logIC_50_=1 corresponds to IC_50_=10μM). In this way, differences between two drug response values are expressed as orders of magnitude in the molar scale.

gdsc_manova_input_w1.csv also contains genetic mutation data for 68 cancer genes (these were selected as the most frequently mutated cancer genes^8^ and their mutational statuses characterise each of the 638 cell lines). For each gene-cell pair, a ‘x::y’ description is provided, where ‘x’ specifies a coding variant and ‘y’ states copy number information from SNP6.0 data. As usual^8^, a gene for which a mutation is not detected in a given cell line is annotated as wild-type (wt). A gene mutation is annotated if a) a protein sequence variant is detected (x≠{wt,na}) or b) a gene deletion/amplication is detected. The latter corresponds to a copy number (cn) range different from the wt value of y=0<cn<8. Furthermore, three genomic translocations were considered (BCR_ABL, MLL_AFF1 and EWS_FLI1) by the GDSC. For each of these gene fusions, cell lines are identified as not-detected fusion or the identified fusion is stated (i.e. wt or mutated with respect to the gene fusion, respectively). The microsatellite instability (msi) status of each cell line is also determined and provided. Further details can be found in the original publication^8^.

### GDSC pharmacotranscriptomic data

These GDSC data sets contain transcriptomic data for 13,321 genes. Gene expression data was generated using Affymetrix Human Genome U219 Array Chip and was normalized with the Robust Multi-Array Average method. The number of cell lines with gene expression data in releases 1 and 5 of the GDSC are 571 and 624, respectively. In terms of data in common, both release contains the expression level of 13,321 genes across 624 cancer cell lines. These genes broke down into 12,644 protein coding genes, 47 pseudogenes, 29 non-coding RNA genes and 601 uncharacterized genes according to the HGNC (HUGO Gene Nomenclature Committee).

### Non-overlapping training and test sets by partitioning data in chronological order

There are 127 drugs in common between both releases. Three drugs are included in the first release only (A-769662, Metformin and BI-D1870), whereas the subsequent release contains 12 additional drugs (TGX221, OSU-03012, LAQ824, GSK-1904529A, CCT007093, EHT 1864, BMS-708163, PF-4708671, JNJ-26854165, TW 37, CCT018159 and AG-014699).

Regarding genomic features, cell lines from both releases have been profiled for 71 common gene alterations in cancer. In addition to the three translocations and msi status, the mutational statuses of 67 genes could be considered (i.e. those for the 68 selected genes in the first release except for the mutational status of the WT1 gene, which was not included in the subsequent release). To ensure that we are using exactly the same drug-gene associations as in the GDSC study, we directly employ the associations and their p-values as downloaded from release 1.

There are two non-overlapping data sets per drug. The training set containing the cell lines tested with the drug and with gene expression data in release 1 (the minimum, average and maximum numbers of cell lines across training data sets are 237, 330 and 467, respectively), along with their IC_50_s for the considered drug. The test set contains the new cell lines tested with the drug and with gene expression data in release 5 (the minimum, average and maximum numbers of cell lines in the test data sets are 42, 171 and 306, respectively). Thus, a total of 254 pharmacotranscriptomic data sets were assembled and analysed for this study.

### Measuring predictive performance of a classifier on a given data set

The pharmacotranscriptomic data for the i^th^ drug (D_i_) can be represented as follow

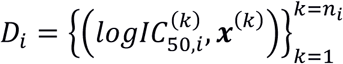

in which, the sensitivity of cancer cell lines against the i^th^ drug has been tested on n_i_ cell lines. **x** is the vector with 13,321 gene expression values. The data can act as training set, cross-validation fold or test set of any of the tested drug.

First, a cell line sensitivity threshold is defined to distinguish between those resistant or sensitive to a given drug. For each drug, we calculated the median of all the logIC_50_ values from training set cell lines and fix it as the threshold. Cell lines with logIC_50_ below the threshold are hence sensitive, while those with logIC_50_ above the threshold are resistant to the drug of interest.

Upon using the model to make predictions in a given data set, two different sets of cell lines will be obtained for each drug: those predicted to be sensitive and those predicted to be resistant. Then, using the threshold for the drug, we can assess classification performance by calculating the number of cell lines in each of these four categories: true positive (TP), true negative (TN), false positive (FP) and false negative (FN). These can be summarised by the Matthews Correlation Coefficient (MCC):

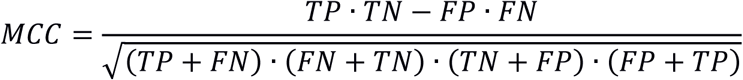

MCC takes values from −1 to 1. An MCC value of 0 means that the tested model has no predictive value, MCC lower than 0 means that the tested model predicts drug sensitivity worse than random and an MCC value equal to 1 indicates that the tested model perfectly predicts the sensitivity of the cell lines against the drug of interest.

In addition to MCC, we also investigated precision (PR), recall (RC) and F1-scores (F1) of the model for each drug to provide a more comprehensive comparison of multi-gene models to single-gene markers. Precision and recall are the two measures of performance for binary classifier, which can be calculated as follow:

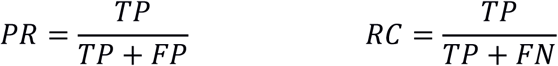

Both metrics can take values from 0 to 1. Precision and recall equal to 0 means that TP = 0, the model fails to identify any cell line sensitive to the drug. By contrast, PR and RC equal to 1 means that FP and FN are equal to 0, respectively. In these cases, either the model does not predict any resistant cell line as sensitive (FP = 0) or it does not misclassify sensitive cell lines as resistant (FN = 0), respectively.

The F1-score is another measure combining PR and RC. F1-score can be computed as

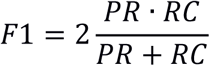

The F1-score is at most 1 (when both PR and RC = 1) and minimum value equal to 0 (RC =0 regardless of the PR value or *vice versa*). High F1 scores mean that both precision and recall are high for the classifier.

### Single-gene markers built from the training data set

We downloaded gdsc_manova_output_w1.csv containing 8701 drug-gene associations with their corresponding p-values computed by MANOVA test. Then, we kept those associations involving the 127 common drugs leading to a set of 8330 drug-gene associations, of which 386 were significant (i.e. p-value smaller than a FDR 20% Benjamini-Hochberg adjusted threshold of 0.00840749). As in previous studies, each statistically significant drug-gene association is regarded as a single-gene marker of *in vitro* drug response.

The best single-gene marker for a drug was identified as its drug-gene association with the lowest p-value. This constitutes a binary classifier with a single independent variable, built using training data alone and fixed at this model selection stage. These drug-lowest p-values were not statistically significant in 15 of the 127 drugs, with the highest of these being P = 0. 0354067. In these cases, we still select them as the best available for these single-gene classifiers. Otherwise, multi-gene markers would be directly better than the single-gene approach for these drugs.

After the model selection step, the single-gene marker for each drug is assessed on the corresponding independent test set. This form of external validation is particularly demanding since the test data is completely separate from training data and constitutes future data from the model training perspective. In 27 drugs, none of the cell lines in the test set harbour the marker mutation and hence TP=FP=0. Therefore, no prediction is provided by these markers and thus MCC and PR are assigned a zero value.

### Multi-gene transcriptomic markers built from the training data set

For each of the 127 drugs, we built a Random Forest (RF) classification model^32^ using exactly the same pharmacological data for training as the corresponding single-gene marker. However, while single-gene markers leverage genomic data, these RF models exploit transcriptomic data instead. All the 13321 gene expression values are used as features (RF_all). Each RF model was built using 1000 trees and the recommended value of its control parameter m_try_ (the square root of the number of considered features, thus m_try_=115 here). All these procedures were implemented in R and executed using Microsoft R Open (MRO) version 3.2.5.

## Supplementary Information

For each analysed drug, results.MNV.RF-CV.RF-TEST.xlsx contains the performances of the best MANOVA-based single-gene marker and RF-based multi-gene marker on the same test set (both methods were in addition trained on the same data set). Furthermore, the 10-fold cross-validated performance of the RF-based multi-gene marker is provided.

## Author contributions

P.J.B. conceived the study, designed its implementation and wrote the manuscript with the help of L.N. L.N. and C.C.D. implemented the software and carried out the numerical experiments. All authors discussed results and commented on the manuscript.

## Acknowledgements

This work has been carried out thanks to the support of the A*MIDEX grant (n° ANR-11-IDEX-0001-02) funded by the French Government «Investissements d’Avenir» programme and the 911 Programme PhD scholarship from Vietnam National International Development.

